# The interplay between motor cost and self-efficacy related to walking across terrain in gaze and walking decisions

**DOI:** 10.1101/2024.08.02.606420

**Authors:** Vinicius da Eira Silva, Daniel S. Marigold

## Abstract

Movement-related decisions, such as where to step and which path to take, happen throughout each day. Shifts in gaze serve to extract task-relevant information necessary to make these decisions. We are only beginning to appreciate the factors that affect this information-seeking gaze behaviour. Here, we aimed to determine how a person’s belief in their ability to perform an action (i.e., self-efficacy) affected gaze and path choice when choosing between walking paths with different terrains and/or lengths (reflecting different energetic costs). We demonstrated that, when manipulated separately (i.e., paths had different lengths or different terrains), participants looked longer and more frequently and chose the paths with either less expected energetic cost or those for which they had a higher self-efficacy rating. When encountering environments where paths differed in both length and terrain, participants directed gaze progressively more to the longer path as the self-efficacy rating of this path increased and the disparity in rating with the shorter (less costly) path grew; they also chose the higher-rated path more frequently regardless of path length. These results provide evidence for a contribution of self-efficacy and energetic cost in guiding gaze and walking decisions. Interestingly, self-efficacy beliefs appear to play a more dominant role in both behaviours.

## INTRODUCTION

Ahead, you are confronted with different walking paths. Each eventually leads to your ultimate destination, but they differ in terms of length, terrain, and/or obstructions. Which one do you choose? Decisions like these are common, and they are likely based on many different considerations. This may include how crowded the path is (Bruneau et al. 2015; Nikmanesh et al. 2024), the energetic cost required to traverse each path (Selinger et al. 2015; Muller et al. 2024), stability constraints associated with each path (Muller et al. 2024), as well as the skill level required to walk across each path without losing balance (da Eira Silva and Marigold 2024). To make appropriate decisions, people can use shifts in gaze to gain timely and accurate information to solve uncertainties about the requirements associated with their action choices (Hayhoe 2017, Domínguez-Zamora et al. 2018; Domínguez-Zamora and Marigold 2021). Here we explore how path length (reflecting energetic cost) and one’s belief in their physical ability affect this information-seeking gaze behaviour and path choice.

Visual information can reveal details about the upcoming terrain and thus, aid in planning foot placement to minimize energy expenditure (Barton et al. 2017; Matthis et al. 2017). Interestingly, people alter their gaze behaviour depending on the cost of stepping or the complexity of the environment ahead (Domínguez-Zamora and Marigold 2019; Matthis et al. 2018; Thomas et al. 2020). For example, when having to expend more energy via greater step widths and redirecting centre of mass to step to a series of targets, there is a delay in gaze shifts away from the current stepping target (Domínguez-Zamora and Marigold 2019). It is unclear whether this link between gaze and motor cost translates to a bias in gazing at the least costly path when deciding between different walking paths.

From an embodied cognition (or choice) perspective, a person perceives action possibilities (or affordances) given the environment and their action capability such that actions and their constraints directly influence decisions (Fajen 2007; Lepora and Pezzulo 2015; Luis-del Campo et al. 2024; van Knobelsdorff et al. 2020). In our path choice example, this implies that someone is less likely to choose a path in which they do not believe capable of crossing without consequence, like when wearing running shoes and presented with icy terrain on a downward slope. People can use the information gathered through gaze to evaluate their belief in their ability to perform an action (i.e., self-efficacy). In a previous study, we demonstrated that people looked more at paths they had a greater level of self-efficacy about walking on, and that behaviour later reflected their path choice (da Eira Silva and Marigold 2024). Thus, how do perceived motor cost and self-efficacy interact to influence gaze behaviour and walking decisions?

To address this question, we asked our participants to walk across one of two paths that we projected on the ground. The paths in each of the environments varied in length and consisted of realistic images of different types of terrain (e.g., dirt, rocks, mud). We recorded participant’s gaze behaviour during the approach phase, since the decision of which path to take happened during that interval. We also had participants rate the confidence in their ability to walk across each terrain type (i.e., self-efficacy rating). For this experiment, we designed three different types of environments. To demonstrate a relationship between self-efficacy and gaze, the first type of environment consisted of paths with the same length but different terrain types. In accordance with our previous study (da Eira Silva and Marigold 2024), people had a greater number of fixations and longer gaze time on the path they rated self-efficacy higher. To demonstrate a relationship between motor cost and gaze, the second type of environment consisted of paths with different lengths but the same terrain types. Our results showed that participants had a greater number of fixations and longer gaze time on the shortest (low energetic cost) path. The third type of environment consisted of different terrain types and different lengths for each path. Thus, paths differed in terms of both motor cost and self-efficacy ratings. Here, we found that the disparity in self-efficacy rating between the high- and low-cost path predicted gaze directed to the high-cost path. Specifically, participants directed gaze more to the high-cost path as the self-efficacy rating of this path increased and the disparity in rating with the low-cost path also increased. This suggests that the brain may rely on self-efficacy to a greater extent than energetic cost when modifying information-seeking gaze behaviour. Together, our results show how the brain considers both energetic cost and self-efficacy about the terrain when making gaze and walking decisions.

## METHODS

### Participants

Sixteen healthy adults participated in this study (3 females and 13 males; mean age = 27.9 ± 7.1 years). Participants did not have any known visual, neurological, muscular, or joint disorder that could affect their walking or gaze behaviour. The Office of Research Ethics at Simon Fraser University approved the study protocol, and participants provided informed written consent prior to the experiment.

### Experimental design

Participants performed a visually guided walking paradigm that required them to walk at a self-selected pace across one of two paths that we projected on the ground (da Eira Silva and Marigold 2024). We created the path images with Photoshop (Adobe Inc., San Jose, CA, USA) and used terrain commonly found on hiking trails, including mud, dirt, and rocks. Each path was approximately 1.2 or 2 m long and 0.55 m wide and joined at the start (see Figure 1). Using images, rather than real terrain, allowed us to maintain better experimental control.

**Figure 1.**
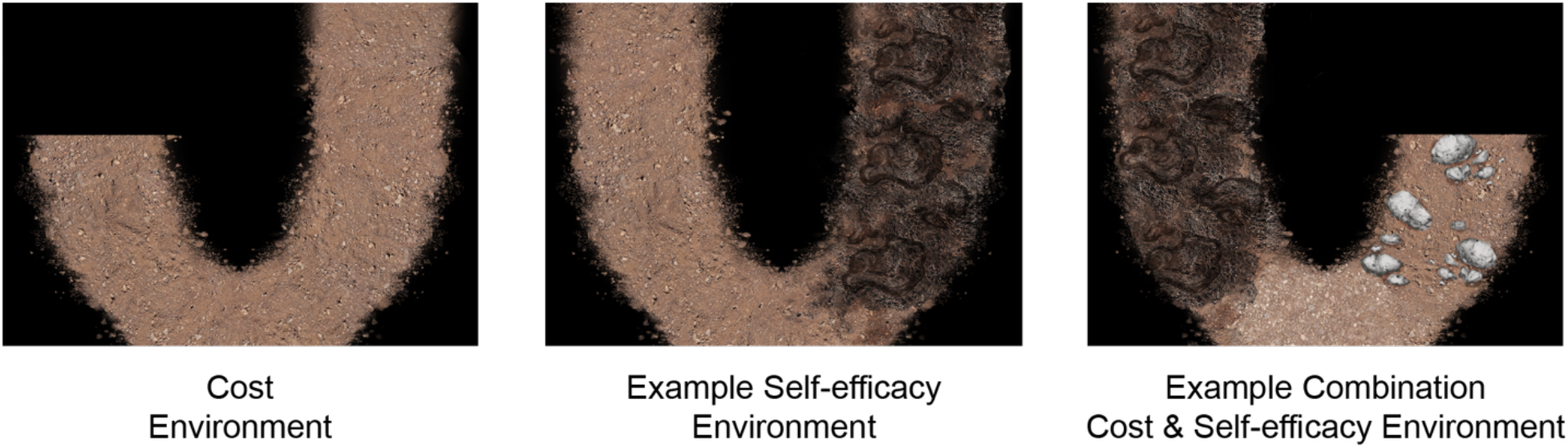
Examples of the environments used in this study. We used one environment to determine the effect of energetic cost, three environments to determine the effect of different self-efficacy ratings, and six environments to determine how people weigh energetic cost and self-efficacy in their gaze and walking decisions. Terrain types include dirt (left panel, both paths), mud (middle panel, right path; right panel, left path), and rocks (right panel, right path). See text for details.

We configured the images of the paths in MATLAB (The MathWorks, Natick, MA) with the Psychophysics Toolbox, version 3 (Brainard 1997), and an LCD projector (Epson PowerLight 5535U, brightness of 5,500 lumens) displayed them on a black uniform mat. To diminish the effect of external references and increase image visibility, participants walked under reduced light conditions (range of 2 to 3 lux, like a moonlit night). We recorded kinematic data at 100 Hz from infrared-emitting position markers placed on the participant’s head and chest and bilaterally on each midfoot (second to third metatarsal head) using two Optotrak Certus motion capture cameras (Northern Digital, Waterloo, ON, Canada) positioned perpendicular to the walking path. We also recorded gaze data at 100 Hz using a high-speed mobile eye tracker (Tobii Pro Glasses 3, Tobii Technology Inc., Reston, VA, USA) mounted on the participant’s head and synchronized with the motion capture system. We calibrated the eye tracker before the walking trials and every 20 trials thereafter based on instructions from the Tobii Pro Glasses user manual.

### Experimental protocol

Before the start of the experimental trials, we showed participants images on the ground of the three types of terrains we used to construct the paths to familiarize them. Subsequently, we used 10 different environments, and participants completed 80 walking trials (eight trials of each environment in random order). Environments had two different paths, where both paths consisted of a single type of terrain. For one environment, participants had to choose between paths with different lengths but the same type of terrain (i.e., dirt). For three environments, participants had to choose between paths with the same length but different types of terrain (i.e., mud vs. dirt, mud vs. rocks, rocks vs. dirt). For six environments, participants had to choose between paths with different lengths and terrains.

For each walking trial, participants started from a standing position approximately 1.5 m from the paths. We projected a fixation cross for one second at the centre of the projection area, approximately 2 m from the starting point, and instructed participants to maintain their gaze on it until the image of the environment appeared. We asked participants to remain stationary and freely visually explore the terrains for two seconds after the fixation cross disappeared until an auditory cue signalled them to begin walking. Participants could also take two steps before they reached the image of the paths. Our choice to allow participants two seconds to explore from a stationary position substituted for the fact that a person would likely see their path choices well in advance (beyond the two steps available in the lab) and could visually explore as they approach from a distance. We asked participants to pretend the terrains were real, to walk at a self-selected speed, to choose the path they would normally take as if outside of the lab, to step where they would normally step if they faced that terrain in real life, and to stop two steps after walking across their chosen path. An experimenter recorded the path that the participant chose each trial.

After completing all experimental walking trials, we had each participant rate the confidence in their ability to walk across each type of terrain (i.e., self-efficacy rating). Specifically, we asked participants: “for each type of terrain, please indicate how confident (or certain) you are of walking across it without losing balance, as though you had to step on it in real life outside. Please use a scale of 1 to 10 (where 1 is not at all confident and 10 is extremely confident).”

### Data and statistical analyses

We filtered kinematic data using a 6-Hz low-pass Butterworth algorithm and used it to calculate the approach phase (defined as the time between the start of the trial and the participant’s first foot contact with the fork in the joined paths). We only analyzed gaze behaviour during the approach phase because we were interested in understanding the decision-making process.

To analyze gaze data, we used GlassesViewer (Niehorster et al. 2020) and a method identical to da Eira Silva and Marigold (2024). We defined fixations as the times during which a target or region on the ground stabilized on the retina. We detected these fixations with the slow-phase classifier described in Hessels et al. (2020) that uses an adaptive velocity threshold based on estimated gaze velocity (classifier parameters: 5000 deg/s start velocity threshold; 50 ms minimum fixation duration; lambda slow/fast separation threshold of 2.5; 8 s moving window). We used the 30 Hz eye tracker video with the gaze location superimposed on the image to verify the presence and location of fixations. To quantify gaze behaviour, we calculated the number of fixations and gaze time (i.e., sum of fixation times) on each terrain, with gaze time normalized by the approach time to control for any differences in gait initiation and speed across trials and participants.

To determine if gaze and path choice are biased to the shorter path, we analyzed the single environment which had different path lengths but identical terrain. Specifically, we compared the number of fixations (or gaze time) during the approach phase on the shorter path with the longer path using separate paired t tests. We predicted a greater number of fixations and gaze times on the shorter path. We used a chi-square test to determine if participants more frequently chose the shorter path.

Da Eira Silva and Marigold (2024) showed that self-efficacy related to walking across terrain affects gaze behaviour and path choice. To demonstrate an effect in this study, we analyzed the three environments which had equal path lengths but differed in the type of terrain. We first confirmed that participants self-efficacy ratings differed between the three different terrains using a mixed-effects model with participant as a random effect and Tukey post hoc tests for a significant effect. For each environment, we compared the number of fixations (or gaze time) during the approach phase between the higher and lower-rated path using separate paired t tests. We predicted a greater number of fixations and longer gaze times on the path with higher a self-efficacy rating. To determine if the difference in self-efficacy ratings between paths predicts gaze behaviour on a path, we performed two linear regressions. In one, we included the proportion of the number of fixations on the left path as the response variable and the difference in self-efficacy ratings between paths (left minus right) as the predictor variable. In the other, we included the proportion of gaze time on the left path as the response variable and the difference in self-efficacy ratings between paths (left minus right) as the predictor variable. We did not include participant as a random effect in these models, because the estimated variance of this effect contributed 0 percent to the total. We used a chi-square test to determine if participants more frequently chose the path that they were more confident in their ability to walk across. For this analysis, we combined data from each of the three environments.

To determine how one’s confidence in their ability to walk across terrain (i.e., self-efficacy) and energetic cost (based on path length) interact to affect gaze behaviour and path choice, we analyzed the six environments where path length and terrain type differed. First, we calculated the proportion of the number of fixations on the longer path and the proportion of gaze time on the longer path for each walking trial. Second, we calculated the difference in self-efficacy rating between the two paths (long path minus short path) for each walking trial. Subsequently, we performed separate linear regressions with either the proportion of the number of fixations on the longer path or the proportion of gaze time on the longer path as the response variable and the difference in self-efficacy rating between paths as the predictor variable. We did not include participant as a random effect in these models, despite the repeated observations, because the estimated variance of this effect contributed 0 percent to the total. We predicted a greater number of fixations and longer gaze time on the more energetically demanding (longer) path when the confidence in their ability to walk across the energetically efficient (shorter) path decreased. We used chi-square tests to determine which path participants were more likely to choose in these environments.

## RESULTS

### Gaze behaviour and path choice are related to the expected energetic cost

In one environment, participants encountered paths with different lengths but identical terrain. To determine if gaze behaviour differed based on the expected energetic cost, we ran separate paired t-tests comparing the number of fixations and gaze time between the shorter path (reflecting less energetic cost) and the longer path (reflecting greater energetic cost). As expected, participants made a greater number of fixations to the low-cost path (6.29 ± 1.26) when compared to the high-cost path (3.48 ± 1.86) (Figure 2A; paired t-test: t_15_=5.09, p=1.33e-4, Cohen’s d =1.27). Participants also had longer gaze time on the low-cost path (0.45 ± 0.12) when compared to the high-cost path (0.22 ± 0.11), the results of which are shown in Figure 2B (paired t-test: t_15_ = 4.20, p = 0.0008, Cohen’s d =1.05). Participants chose the shorter path 88% of the time (χ^2^: p = 3e-14). These results suggest that gaze behaviour and path choice are biased to the path requiring the least energetic cost.

**Figure 2.**
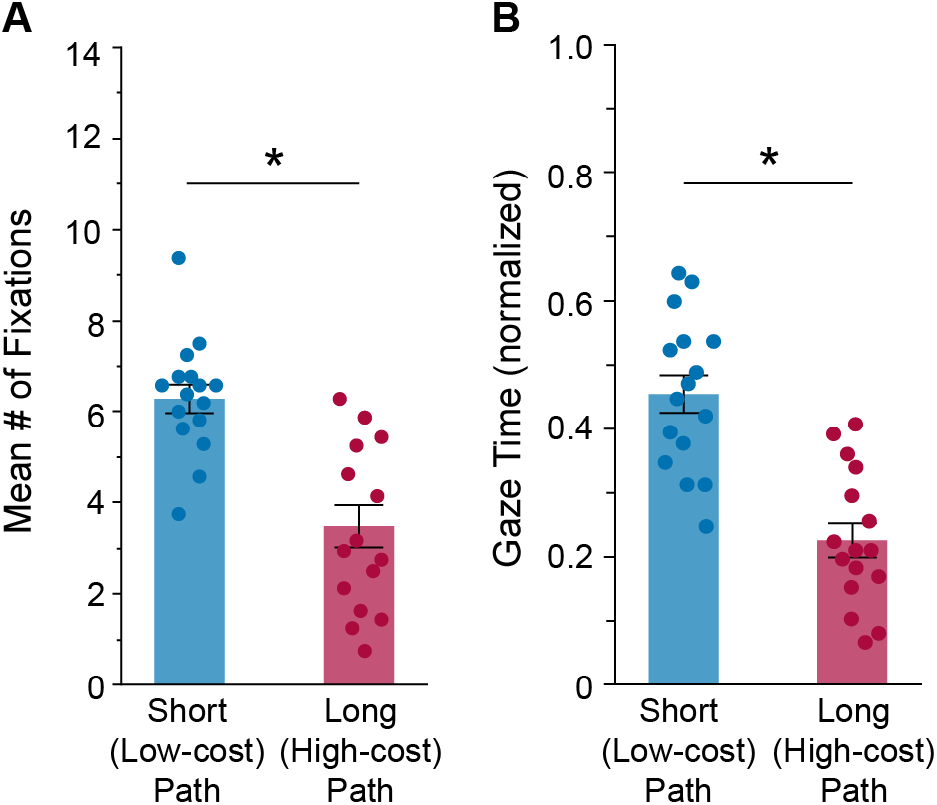
How energetic cost (different path lengths) relates to gaze behaviour during the approach (decision-making) phase. (A) Group mean ± SE number of fixations on the two paths. (B) Group mean ± SE gaze time on the two paths. Mean individual participant (*n* = 16) data values are superimposed. Asterisk indicates a statistically significant difference between the paths (p < 0.05).

### Gaze behaviour and path choice are also related to self-efficacy beliefs

In three environments, participants encountered paths of identical length but that differed in terrain. This allowed us to isolate any effect of self-efficacy. Participant’s self-efficacy ratings differed significantly depending on the terrain types (F_2,30_ = 137.9, p = 7.53e-16). Specifically, post hoc tests revealed a significant difference in ratings between the dirt terrain (9.8 ± 0.4), rocky terrain (8.1 ± 1.0), and mud terrain (5.1 ± 1.1).

To determine if gaze behaviour differed based on self-efficacy ratings, we ran separate paired t-tests comparing the number of fixations and gaze time between the higher and lower rated path for each environment. As illustrated in Figure 3A, we found that participants had a greater number of fixations on the higher-rated self-efficacy path compared to the lower-rated path for the mud vs. dirt environment (t_15_ = -6.02, p = 2.37e-5, Cohen’s d = 1.50), mud vs. rocks environment (t_15_ = - 9.10, p = 1.7e-7, Cohen’s d = 2.28), and rocks vs. dirt environment (t_15_ = -3.88, p = 0.0015, Cohen’s d = 0.97). Also shown in Figure 3B, we found that participants had greater gaze time on the higher-rated path compared to the lower-rated path for the mud vs. dirt environment (t_15_ = -5.15, p = 1.19e-4, Cohen’s d = 1.29), mud vs. rocks environment (t_15_ = -7.83, p = 1.12e-6, Cohen’s d = 1.96), and rocks vs. dirt environment (t_15_ = -5.08, p = 0.0001, Cohen’s d = 1.27).

**Figure 3.**
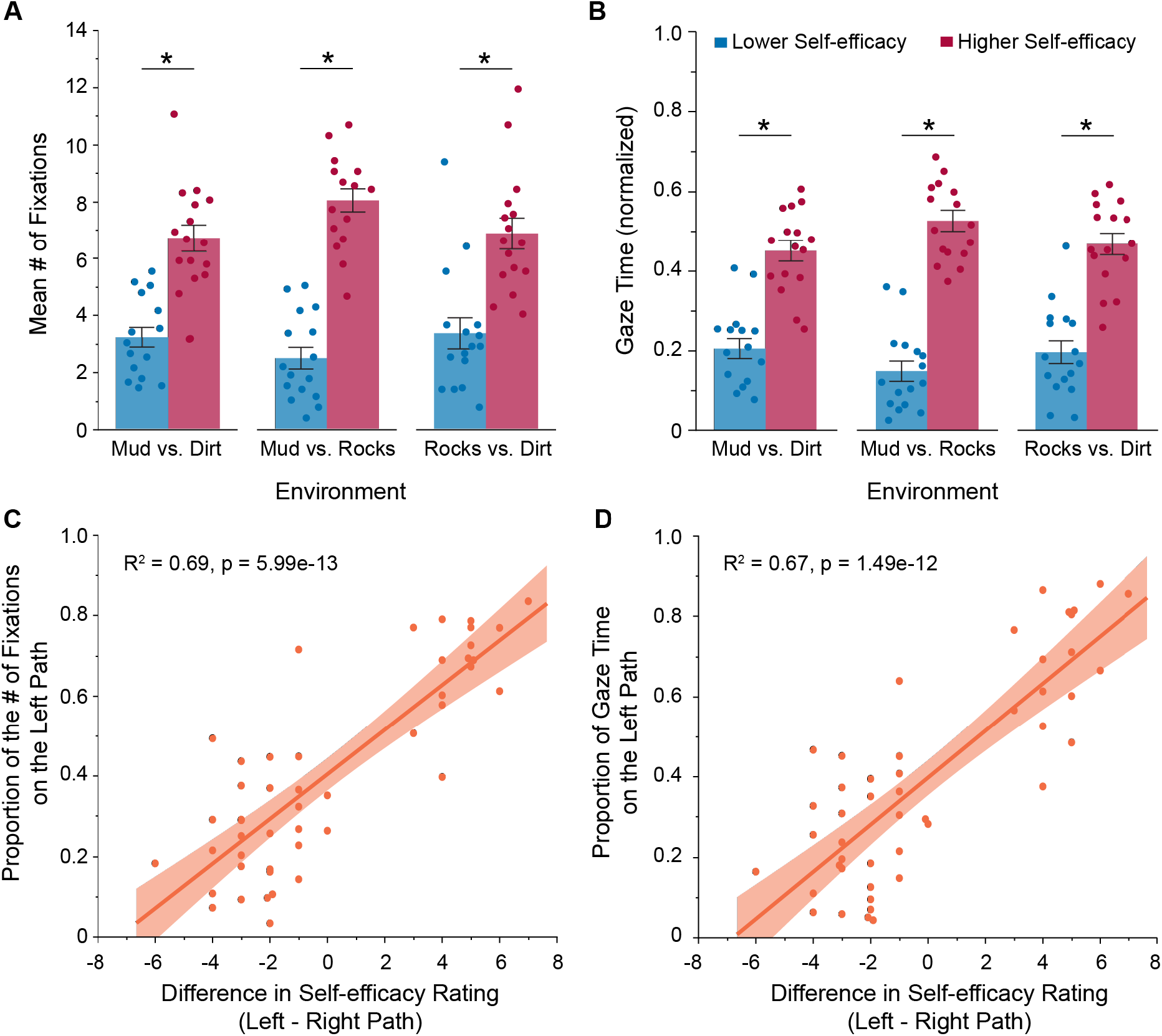
How self-efficacy related to walking across a path influences gaze behaviour during the approach (decision-making) phase. (A) Group mean ± SE number of fixations on the two paths for the three environments where terrain differed but length did not. (B) Group mean ± SE gaze time on the two paths for the three environments where terrain differed but length did not. Individual participant (*n* = 16) data values are superimposed. (C) Scatterplot of the proportion of the number of fixations on the left path versus the difference in self-efficacy rating between the two paths. (D) Scatterplot of the proportion of gaze time on the left path versus the difference in self-efficacy rating between the two paths. In each scatterplot, solid lines show the linear fits obtained from the models and shaded regions represent the 95% confidence intervals. Asterisk indicates a statistically significant difference between the paths (p < 0.05).

Since participants chose the path that they rated higher in terms of self-efficacy 96% of the time (three environments combined; χ^2^: p = 4e-20), it is possible that the gaze behaviour during the approach phase reflects, at least in part, the path they intend to take rather than an effect of self-efficacy on gaze. Thus, we determined how gaze varied on one path based on the difference in self-efficacy ratings between paths, independent of path choice. If gaze simply reflects the intended path, it is reasonable to expect a non-significant (flat) slope in the regression. This was not the case. As illustrated in Figure 3C,D, the difference in self-efficacy between the two paths predicted the number of fixations (R^2^ = 0.69, p = 5.99e-13) and gaze time (R^2^ = 0.67, p = 1.49e-12) on the left path. These results suggest that gaze behaviour and path choice are biased to the path that participants are more confident in their ability to walk across.

### People prioritize self-efficacy over energetic cost when deciding where to direct gaze and walk

In six environments, participants encountered paths that differed in both length and terrain. This allowed us to determine how people trade-off expected energetic cost for confidence in their ability to walk across a certain terrain in their decision of where to look and where to walk. We calculated the difference in self-efficacy ratings between paths (long, higher-cost path minus short, lower-cost path) and used it as the predictor variable in separate linear regressions. For the response variable, we used either the proportion of the number of fixations or gaze time on the long (higher-cost) path. As illustrated in Figure 4, we found that the difference in self-efficacy between paths predicted gaze behaviour (number of fixations: R^2^ = 0.46, p = 3.01e-13; gaze time: R^2^ = 0.46, p = 3.43e-13). Participants increased fixations and spent more time looking at the longer (higher-cost) path as the confidence in their ability to walk across this path increased relative to the short (lower-cost) path.

**Figure 4.**
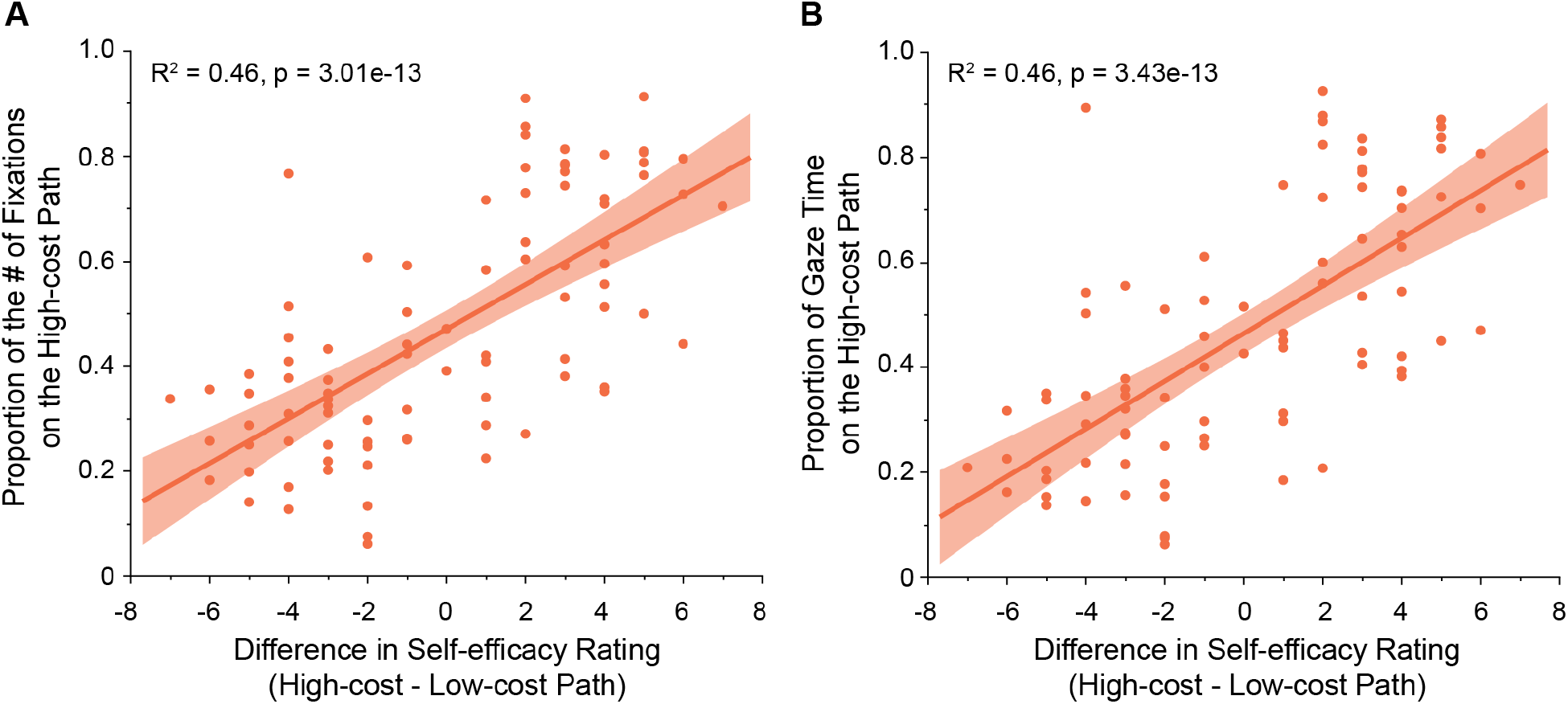
Trade-off between energetic cost and self-efficacy related to walking across terrain in the decision of where to direct gaze during the approach (decision-making) phase. (A) Scatterplot of the proportion of the number of fixations on the high-cost path versus the difference in self-efficacy rating between the two paths. (B) Scatterplot of the proportion of gaze time on the high-cost path versus the difference in self-efficacy rating between the two paths. In each scatterplot, solid lines show the linear fits obtained from the models and shaded regions represent the 95% confidence intervals.

Participants preferred to take the higher self-efficacy-rated path regardless of path length. Specifically, participants chose the shorter, higher self-efficacy-rated path 47.7 ± 4.9 % of the time, the longer, higher-self-efficacy-rated path 37.3 ± 14.7 % of the time, the shorter, lower self-efficacy-rated path 12.6 ± 13.6 % of the time, and the longer, lower self-efficacy-rated path 2.4 ± 4.3 % of the time. When considering self-efficacy ratings, participants chose the higher-rated path 85% of the time (χ^2^: p = 3e-12). When considering energetic cost, participants chose the shorter (lower-cost) path only 60% of the time (χ^2^: p = 0.046).

## DISCUSSION

To make a decision, people seek out and weigh incoming information about available options based on a variety of considerations. During walking, these considerations are particularly important given that an incorrect or suboptimal decision can have meaningful consequences. For instance, people tend to care about saving energy as they walk (Selinger et al. 2015), but they also prefer paths for which they rate themselves has having higher self-efficacy in crossing (da Eira Silva and Marigold 2024). Here, we aimed to determine how self-efficacy affected gaze and path choice when choosing between paths with different energetic costs and/or terrains. We used a forced-choice paradigm, where we manipulated the types of terrain and/or lengths in each path. We demonstrated that, when manipulated separately (i.e., paths had different lengths or different terrains), participants looked longer and more frequently and chose the paths with less expected energetic cost or those for which they had a higher level of self-efficacy rating. When facing environments where paths differed in both lengths and terrains, participants directed gaze progressively more to the longer path as the self-efficacy rating of this path increased and the disparity in rating with the shorter path grew; they also chose the path in which they gave a higher self-efficacy rating more frequently regardless of path length. These results suggest a potential role for self-efficacy in gaze and walking decisions. In the following paragraphs, we situate our results within a proposed framework for gaze and walking decisions and discuss limitations of our work.

There are at least two decision algorithms engaged in our walking task: where and when to direct gaze and which path to walk. The main role of gaze in a goal-directed action is to obtain task-relevant information (or reduce uncertainty) about the environment (Domínguez-Zamora et al. 2018; Domínguez-Zamora and Marigold 2021; Gottlieb et al. 2014; Hayhoe 2017; Tong et al. 2017). Given the number of possible locations in the environment to look at, and the limited time available when walking, the brain must determine how best to extract information applicable to the action. There are likely several factors that can influence this process. We propose a framework where self-efficacy beliefs and motor cost modify information-seeking gaze behaviour. This conceptual framework is illustrated in Figure 5.

**Figure 5.**
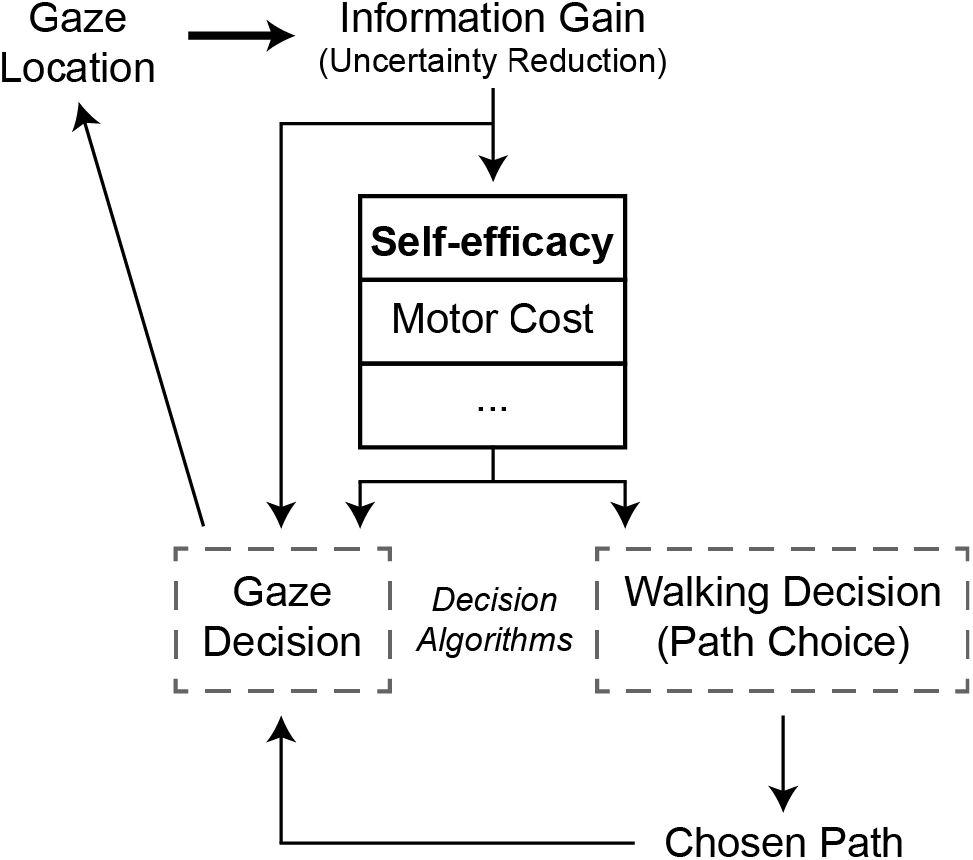
Proposed conceptual framework for gaze and walking decisions. The main role of gaze is to provide information (or reduce uncertainty) about the environment. We argue that self-efficacy related to walking across terrain and motor cost serve to modify information-seeking gaze behaviour in addition to path choice.

Given that people typically look where they intend to walk (Domínguez-Zamora and Marigold 2021; Marigold and Patla 2007), it is possible that gaze behaviour simply reflects the decision of which path to take and thus, self-efficacy indirectly affects gaze. However, the results of the current study, and our previous work (da Eira Silva and Marigold 2024), support the involvement of self-efficacy beliefs in the gaze decision. This is evident in Figures 3 and 4, which show that gaze metrics on one path vary with the difference in self-efficacy between paths across different environments. Specifically, we found that people are more likely to focus gaze on (or exploit) the path that they are more confident in their ability to manage when there is a clear difference between options. However, they are more likely to explore both options when the confidence in their ability to manage each one is similar. In this latter situation, when the difference in self-efficacy ratings was between -1 and 1, we show that the proportion of fixations or gaze time on one path varies between 40 and 50% based on the fit of the regression line (see Figure 4). Together, these results suggest that gaze is not merely a reflection of the walking decision.

Although we found that gaze was biased toward the higher self-efficacy-rated path, rather than to the shorter (less costly) path, past research has demonstrated a role for motor cost in affecting gaze behaviour. For instance, the shift in gaze from the current to subsequent step location is delayed when stepping wider, a movement that requires greater energy expenditure from increased muscle activity (Domínguez-Zamora and Marigold 2019). Furthermore, how complex (or how costly) the terrain is to walk across regulates how many steps ahead gaze is directed (Matthis et al. 2018). Thus, we believe it is appropriate to include motor cost as an input to the gaze decision algorithm of our framework.

Based on the visual information extracted from the environment, one can make an informed decision about where to walk. Here we demonstrated that self-efficacy beliefs strongly influenced the walking decision, like in our previous work (da Eira Silva and Marigold 2024). We also found that motor cost had an impact on this decision, but clearly not to the same extent as self-efficacy. However, others have demonstrated that motor cost affects the selection of where to step and route planning over rough terrain (Domínguez-Zamora and Marigold 2021; Moraes 2014; Muller et al. 2024), as well as the decision of which direction to walk around an obstacle (Grießbach et al. 2022); though people do not necessarily choose the most energetically efficient path up a hill (Linkenauger et al. 2019). Thus, in our framework, both self-efficacy beliefs and motor cost also contribute to the walking decision.

Are there other explanations for the observed gaze and walking decisions beyond self-efficacy? It is possible that the perceived dynamic stability costs associated with the path’s terrain influenced behaviour. Under this view, participants chose to look and walk across the longer path in the mixed environments—where terrains and path lengths both differed—because they perceived it to have a lower stability cost based on past experiences with similar terrain, rather than because the longer path may have had higher self-efficacy ratings. Self-efficacy beliefs are acquired through past performance, vicarious experiences, verbal persuasion (or motivation), and arousal (Bandura 1977, 1997). Stability costs are likely to at least partially influence the first and last of these sources. The effect of stability cost on self-efficacy would depend on the past performance itself and the frequency of encountering the type of terrain. For instance, repeated successful negotiation of rocky or muddy terrain may make a person believe that they are capable of walking across them again without issue, leading to higher self-efficacy beliefs. In contrast, previously tripping or falling when encountering this terrain may lead to relatively lower self-efficacy beliefs. Thus, how a participant perceived the stability costs of the terrain may have contributed to their self-efficacy rating and behaviour.

If the perceived cost to maintain dynamic stability contributes to the choice of looking at and walking across the longer path, then a possible implication is that this cost may outweigh the cost associated with walking a longer distance. In other words, participants could have perceived the overall cost for the shorter path (with mud or rocks, for example) as greater than the overall cost of the longer path (with dirt, for example). As an estimate of energy expenditure, we assumed a constant walking velocity and multiplied path length by a terrain coefficient (or terrain factor) to account for the cost of the type of terrain. For the terrain factor, we used known differences in energy expenditure (Kim et al. 2023; Voloshina et al. 2013) or established terrain factors (Richmond et al. 2015) based on terrain like those used in our experiment, including a dirt path, flat terrain with obstacles (or mixed terrain), uneven terrain, and mud. Despite factoring in the cost of different terrain, the longer path was still more costly overall compared to the shorter path in each of our environments. Furthermore, although we instructed participants to treat the terrain images as though they were real, there were no actual stability costs in our task; the only true cost related to walking distance. Taken together, we argue that a person’s self-efficacy beliefs likely drove our participant’s behaviour.

Overall, we show how self-efficacy and motor cost are used in the process that guides gaze and walking decisions. One’s confidence in one’s ability to perform an action (i.e., self-efficacy) appears to play a particularly important role in decision-making. This idea may relate to a desire to prevent injury, and from an evolutionary point of view, to ensure survival. How self-efficacy interacts with other potential inputs of decision algorithms is unclear. Nonetheless, our work has broad implications for how we gather information and make decisions and provides important insight to advance current models of gaze behaviour for goal-directed, natural actions (Hayhoe 2017; Sprague and Ballard 2003).

## ACKNOWLEDGEMENTS

The authors thank Laura Gimenes for help with terrain illustrations.

